# A genetically engineered phage-based nanomaterial for detecting bacteria with magnetic resonance imaging

**DOI:** 10.1101/2022.06.07.495091

**Authors:** Raymond E. Borg, Harun F. Ozbakir, Binzhi Xu, Eugene Li, Xiwen Fang, Huan Peng, Irene A. Chen, Arnab Mukherjee

**Affiliations:** Department of Chemistry, University of California, Los Angeles, CA 90095, USA; Department of Chemical Engineering, University of California, Los Angeles, CA 90095, USA; Biomolecular Science and Engineering, University of California, Los Angeles, CA 90095, USA; Biological Engineering, University of California, Los Angeles, CA 90095, USA; Neuroscience Research Institute, University of California, Santa Barbara, CA 93106, USA; Department of Chemical and Biomolecular Engineering, University of California, Los Angeles, CA 90095, USA

## Abstract

The ability to noninvasively detect bacteria at any depth inside opaque tissues has important applications ranging from infection diagnostics to tracking therapeutic microbes in their mammalian host. Current examples of probes for detecting bacteria with strain-type specificity are largely based on optical dyes, which cannot be used to examine bacteria in deep tissues due to the physical limitation of light scattering. Here, we describe a new biomolecular probe for visualizing bacteria in a cell-type specific fashion using magnetic resonance imaging (MRI). The probe is based on a peptide that selectively binds manganese and is attached in high numbers to the capsid of filamentous phage. By genetically engineering phage particles to display this peptide, we are able to bring manganese ions to specific bacterial cells targeted by the phage, thereby producing MRI contrast. We show that this approach allows MRI-based detection of targeted *E. coli* strains while discriminating against non-target bacteria as well as mammalian cells. By engineering the phage coat to display a protein that targets cell surface receptors in *V. cholerae*, we further show that this approach can be applied to image other bacterial targets with MRI. Finally, as a preliminary example of *in vivo* applicability, we demonstrate MR imaging of phage-labeled *V. cholerae* cells implanted subcutaneously in mice. The nanomaterial developed here thus represents a path towards noninvasive detection and tracking of bacteria by combining the programmability of phage architecture with the ability to produce three-dimensional images of biological structures at any arbitrary depth with MRI.

## INTRODUCTION

The ability to detect specific bacterial strains in deep-seated tissue has important clinical applications, ranging from accurate noninvasive diagnostics and treatment monitoring to tracking bacteria-based live biotherapeutic products *in vivo*^1^. In a preclinical setting, targeted imaging of bacteria in vertebrate models could aid basic research on infectious disease mechanisms and host-microbe interactions as well as facilitate the development of bacterial formulations for treating diseases like cancer and gastrointestinal inflammation^2,3^. Given that light scattering limits the use of optical imaging to shallow (sub-millimeter) depths, tissue-penetrant modalities are required to detect bacteria in deep tissues. To this end, probes based on positron emission tomography (PET) represent the current gold standard for deep-tissue imaging of bacteria using radioactive tracers conjugated to ligands that broadly target bacteria, for example antibodies, antibiotics, antimicrobial peptides, and cell-internalizable sugars^4–9^. While PET achieves high detection sensitivity, the spatial resolution is limited (millimeter-scale in rodents); and potentially harmful ionizing radiation is required to penetrate tissues.

In contrast to PET, magnetic resonance imaging (MRI) can produce 3-dimensional images with high resolution (∼ 100 μm) and excellent soft tissue contrast, while avoiding exposure to ionizing radiation. In clinical practice, these capabilities are already leveraged to provide diagnostic information on soft tissue infections. However, the basis for detecting infections with MRI is largely nonspecific, relying on changes in tissue structure at the infected site; and only a handful of bacterially-targeted MRI probes have been reported. The first example of such a probe coupled a paramagnetic Gd(III) chelate with cationic zinc complexes that bind to the negatively charged phospholipids on the bacterial membrane, thereby allowing bacteria to be specifically monitored with (T_1_ weighted) MRI^10^. More recently, broad-spectrum antibiotics like neomycin and vancomycin have been employed as targeting groups to localize Gd(III) based contrast agents to bacterial cells^11–13^. Bacterial detection has also been attempted by filling cells with iron-oxide nanoparticles^14–16^, which produce T_2_ weighted MRI contrast. While the aforementioned probes can provide useful information on general bacteria classes (*e*.*g*., gram positive), they lack well defined cell-type specificity. Furthermore, iron oxide based probes give rise to negative contrast, *i*.*e*., signal darkening, which can be hard to differentiate from non-specific sources of signal dropout such as air-tissue interfaces, abscesses, and microhemorrhage. Finally, probes that make use of antimicrobial ligands as the targeting agent could exhibit bactericidal activity, which may not always coincide with the intended goal of an application, for example tracking bacterial therapeutics *in vivo*.

Here we investigate an alternative biomolecular scaffold for imaging bacterial cells with high specificity using the non-lytic, filamentous bacteriophage, M13. This approach leverages the malleability of the phage capsid to attach large numbers (∼ 2700 per phage) of imaging probes on the major coat protein (known as pVIII) (**Figure 1A**). These phages can then be targeted to defined cell-types by engineering a second coat protein (for example, pIII) to express proteins that bind to specific receptors in in the target microorganism. Phages have been extensively engineered to detect bacteria with techniques like fluorescence, absorbance, and impedance spectroscopy as well as PET and near infrared imaging (Table S2)^17–21^. However, to our knowledge, the application of phages for imaging bacteria with MRI has not been reported. Therefore, with the goal of combining cell-type targetability of phage with the high resolution, nonionizing, and deep tissue-penetrant capabilities of MRI, we engineered M13 to develop a paramagnetic nanomaterial for imaging bacteria with MRI. We characterize the engineered phages by relaxometric measurements and demonstrate their utility for visualizing F-pilus expressing *E. coli* cells with high specificity. Furthermore, we modified phage tropism by engineering the pIII coat protein to target a pathogenic bacterium, *Vibrio cholerae*, and applied the ensuing sensors to image subcutaneously implanted bacteria in a mouse model.

**Figure 1.**
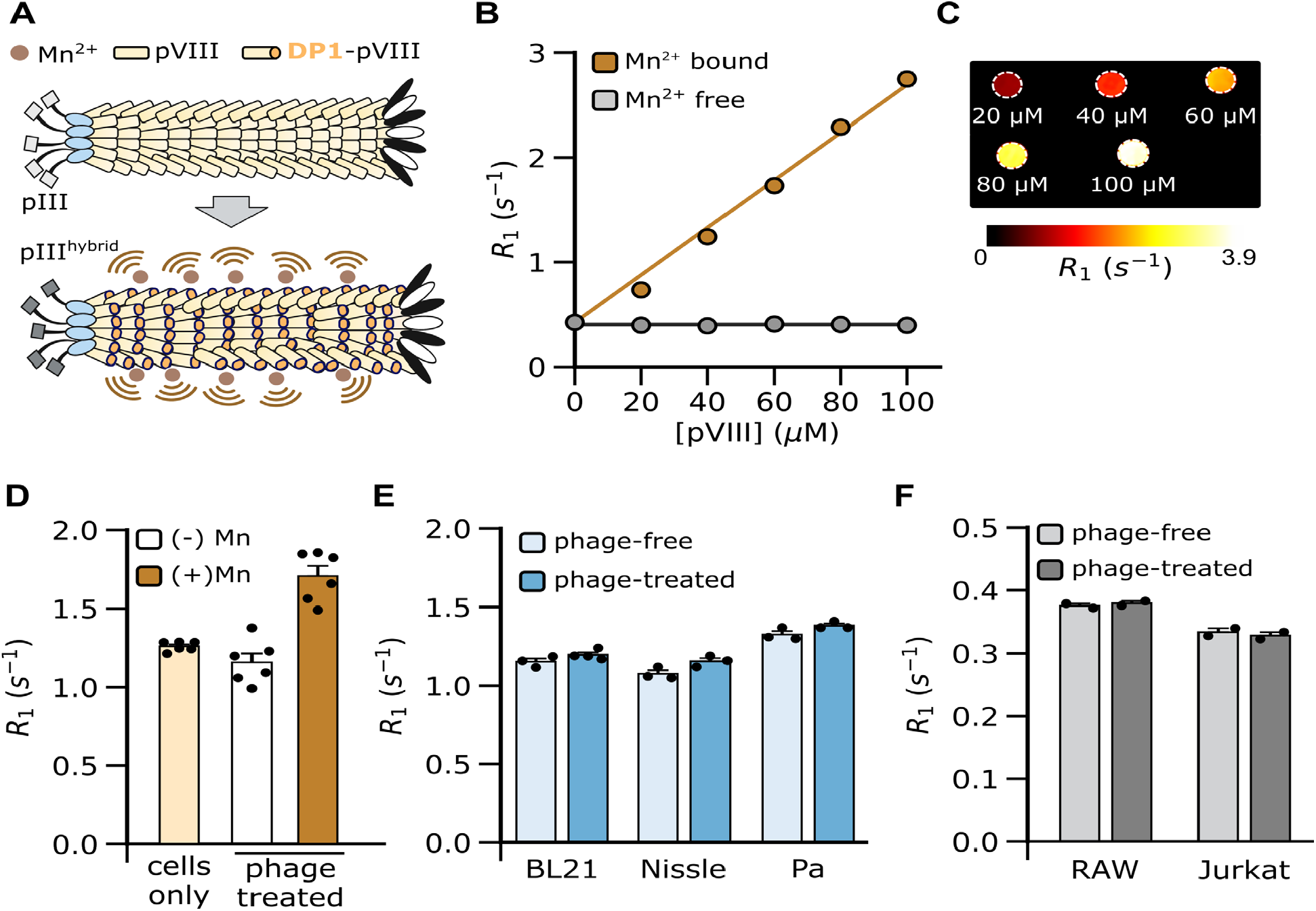
Bacterial detection with phage-based MRI sensors. **(A)** Overview of imaging approach. DP1 is genetically attached to the exposed N-terminus of pVIII, allowing the phage coat to multivalently bind manganese. The phage is then used to bring manganese ions close to bacterial cells targeted by the native pIII protein or engineered hybrid variants (denoted as pIII^hybrid^), thereby enabling detection with MRI. **(B)** T_1_ relaxation rate (R_1_) of manganese-bound and manganese-free DP1-phage measured in HEPES buffer. Error-bars are smaller than the marker size. **(C)** R_1_ maps of manganese-bound DP1-phage at phage titers corresponding to pVIII concentrations in the 0-100 uM range. R_1_ maps were generated by manually masking the region of interest (ROI) and fitting the signal intensity in each voxel within this ROI to an exponential equation describing T_1_ relaxation. The images were smoothed with a 2-pixel median filter and displayed on a linear 8-bit heatmap by thresholding at minimum and maximum values of R_1_ as depicted in the adjoining color bar. **(D)** R_1_ of phage-free and phage-treated cultures of *E. coli* (F^+^) cells. A significant increase in R_1_ relative to cells-only baseline (*p = 4*.*9 × 10*^*-5*^, *N = 6*) is observed after incubating cells with manganese-loaded (but not manganese-free) DP1-phage. **(E)** R_1_ of phage-free and phage-treated cultures of non-target bacterial strains. BL21 - *E. coli* BL21, Nissle - *E. coli* Nissle, and Pa - *Pseudomonas aeruginosa*; (F) R_1_ of phage-free and phage-treated mammalian cell lines: Jurkat (T cell) and RAW264.7 (macrophage). R_1_ values were comparable for the non-target cells in both phage-treated and phage-free conditions. MRI measurements were performed at 7 Tesla. Error bars depict standard error from N ≥ 3 replicates.

## RESULTS AND DISCUSSION

### Preparation of manganese-loaded M13 phage

Our basis for engineering an MRI-detectable phage involves a manganese-binding peptide, DP1 (**Table S1**), first identified in the radiation-resistant extremophile, *D. radiodurans*^22–24^. We hypothesized that phages could be made MRI-visible by genetically modifying the phage coat to display DP1, which would then allow the phage to specifically bind manganese, a well-established T_1_ weighted MRI contrast agent. The resulting phage could then be directed to bind specific bacterial species based on the composition of the pIII coat protein, thereby bringing manganese ions to the target bacterial cells for imaging with MRI (**Figure 1A**). We chose the DP1 peptide because of its natural affinity towards Mn^2+^ (K_d_ ∼ 10 μM^23^) as well as its small size (10 amino acids), which makes it convenient for forming genetic fusions with the pVIII major coat protein (of which there are ∼ 2700 copies per phage), thereby enabling the phage capsid to bind multivalently with manganese. To stay within the size limit tolerated by pVIII fusions (typically 6-8 amino acids^25^), we trimmed DP1 down to 8 amino acids without loss of manganese binding, as determined by a fluorescence assay (**Figure S1**). We cloned the truncated DP1 at the exposed N-terminus of pVIII in a packaging plasmid harboring all nine phage-forming proteins, and the correct clone was verified by DNA sequencing. We produced phage particles by transforming *E. coli* with the packaging plasmid along with a second plasmid containing an M13 origin of replication. DP1-phages were precipitated by treating culture supernatants with NaCl and poly(ethylene) glycol (PEG), and phage formation was verified with absorbance spectroscopy, Western blotting, and size-exclusion chromatography (**Figure S2A-B**).

Phages were loaded with manganese by incubating with MnCl_2_ and purified by three additional rounds of PEG-NaCl precipitation to remove unbound manganese ions. Under these conditions, we were able to achieve a manganese loading of 47.3 ± 0.9% (relative to pVIII concentration), as measured with inductively coupled plasma-mass spectrometry (ICP-MS) (**Figure S2C**).

### Relaxometric characterization of DP1- phage

Next, we tested the ability of the manganese-bound phage to enhance the baseline T_1_ relaxation rate (R_1_) of water protons. Manganese-loaded DP1-phage was found to increase R_1_ in proportion to phage concentration, achieving a peak increase of 549.7 ± 14 % at a phage titer that corresponds to 100 μM pVIII (**Figures 1B-C**). In contrast, R_1_ was unchanged, *i*.*e*., similar to aqueous buffer, for DP1-phage that was not pre-incubated with manganese (**Figure 1B**). Furthermore, the molar relaxivity (defined as the slope of R_1_ *vs*. [pVIII]) of phage was unaffected (*p > 0*.*6, N = 4*) by incubation with physiologically relevant concentrations of common divalent metals such as Zn^2+^, Ca^2+^, and Mg^2+^ indicating the selectivity of DP1 towards manganese (**Figure S3A**). When assayed in mouse serum and cell culture medium, DP1- phage showed a moderate decrease in molar relaxivity compared to values measured in aqueous buffer (**Figure S3B**). Given that these physiological formulations contain a complex mixture of cofactors, metal salts, proteins, and other macromolecules, it is possible that some components could interfere with the interaction between DP1- phage and manganese ions, thereby lowering phage relaxivity.

### MRI detection of a specific *E. coli* strain

To assess whether the paramagnetic properties of DP1-phage could be used to detect bacteria with MRI, we leveraged the natural ability of the pIII coat protein to attach to *E. coli* (F^+^) strains by binding to the F-pilus receptor with picomolar affinity^26^. Overnight cultures of an *E. coli* (F^+^) strain were incubated with DP1-phage (pre-loaded with manganese) at a phage-to-cell stoichiometry of 50:1 and washed twice to remove unbound virions. When assessed with MRI, the relaxation rate of phage-treated *E. coli* (F^+^) cells was found to be 35 ± 1.4 % faster relative to cells that had not been incubated with phage (*p = 4*.*9 × 10*^*-5*^, *N = 6*) (**Figures 1D, S4**). No change in relaxation rate (*p = 0*.*11, N = 6*) was obtained when cells were incubated with manganese-free phage (**Figure 1D**), indicating that DP1-phage by itself, *i*.*e*., without manganese, does not accelerate T_1_ relaxation, which is consistent with the diamagnetic nature of manganese-free phage suspensions (**Figure 1B**). To test whether detection was specific to *E. coli* (F^+^) cells, we examined the ability of DP1-phage to generate MRI contrast following incubation with representative bacterial strains lacking the F-pilus, including *E. coli* BL21, *E. coli* Nissle, and *P. aeruginosa*; as well as two mammalian cell lines: Jurkat (T cell line) and RAW264.7 cells (macrophage cell line). No substantial change in R_1_ was observed with any of these cell-types (**Figures 1E-F, S4**), confirming specificity of the DP1-phage towards F-pilus-expressing *E. coli* cells.

### Engineering DP1-phage to detect *V. cholerae in vitro*

Having established the basic capabilities of DP1-phage for detecting bacteria with MRI, we next engineered a phage to recognize *V. cholerae*, a pathogenic bacterium that continues to pose a major threat to public health, disproportionately affecting populations with limited access to clean water, sanitation, and healthcare facilities. To target *V. cholerae*, we substituted the F-pilus-binding residues of the pIII coat protein (amino acids 18-236) with corresponding residues from the minor coat protein of phage CTXΦ, which binds a cell surface receptor (toxin-coregulated pilus^27^) in *V. cholerae*, resulting in a hybrid phage DP1-CTX. This approach has been previously shown to direct a chimeric phage to bind *V. cholerae* cells^28,29^; however, its application for imaging cells with MRI (or any tissue-penetrant modality) has not been reported. The dual-functionalized DP1-CTX phage particles were purified, loaded with manganese as before, and shown to generate comparable T_1_ relaxivity as the DP1-phage (**Figure 2A, S5A**). To assess whether DP1-CTX phage could be used to target *V. cholerae*, we treated overnight cultures with purified phage particles and imaged the cells at 7 Tesla as described earlier. Phage-labeled *V. cholerae* showed a 61 ± 4 % increase in their relaxation rate relative to phage-free cultures (*p = 10*^*-5*^, *N = 6*). As expected, no change in R_1_ was detected when *V. cholerae* cells were incubated with DP1-CTX phage not loaded with Mn^2+^ (*p = 0*.*4, N = 3*) (**Figure 2B**). To determine whether detection was specific to *V. cholerae*, we characterized relaxation rates in *E. coli* BL21, *E. coli* Nissle, and *P. aeruginosa* cultures incubated with DP1- CTX phage following exactly the same procedures as described above. In all cases, changes in R_1_ were markedly smaller, confirming specificity of DP1-CTX phage for *V. cholerae* (**Figures 2C, S5B**).

**Figure 2.**
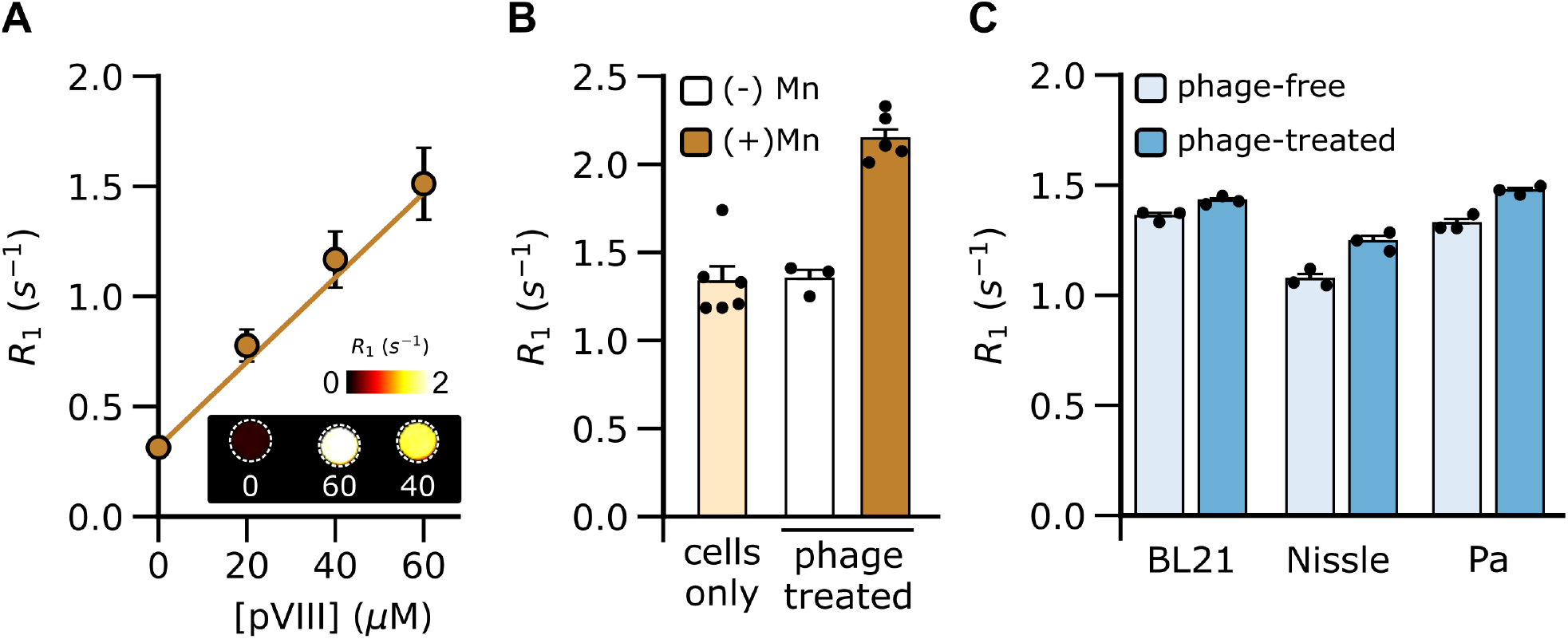
MRI-based detection of *V. cholerae* with DP1-CTX phage. **(A)** T_1_ relaxation rate (R_1_) of manganese-bound and manganese-free DP1-CTX phage measured in HEPES buffer. Inset: R_1_ maps of manganese-bound DP1-CTX phage corresponding to pVIII concentrations from 0-60 μM. R_1_ maps were generated by manually masking the region of interest (ROI) and fitting the signal intensity in each voxel to an exponential equation describing T_1_ relaxation. The images were smoothed with a 2-pixel median filter and displayed on a linear 8-bit heatmap by thresholding at 0 and 2 s^-1^ as depicted in the adjoining color bar. **(B)** R_1_ of phage-free and phage-treated *V. cholerae* cells. A significant increase in R_1_ relative to cells-only baseline (*p = 2 × 10*^*-5*^, *N = 6*) is observed after incubating cells with manganese-loaded (but not manganese-free) DP1-CTX phage. **(C)** R_1_ of DP1-CTX phage-free and phage-treated cultures of non-target bacteria. BL21 - *E. coli* BL21, Nissle - *E. coli* Nissle, and Pa - *Pseudomonas aeruginosa*. All MRI measurements were performed at 7 Tesla. Error bars represent standard error of mean from N ≥ 3 replicates.

### MRI detection of *V. cholerae in vivo*

Finally, to provide a preliminary assessment of the capabilities of DP1-CTX phage to track bacteria deployed *in vivo*, we labeled *V. cholerae* cells with phage particles as described above and injected the labeled cells in the hind flanks of mice. The contralateral flank was implanted with unlabeled, *i*.*e*., phage-free, *V. cholerae* to provide a well-matched control in the same animal. When imaged at 7 Tesla, the T_1_ relaxation rate was found to be 54 ± 6 % faster in the flanks harboring phage-labeled bacteria compared to the flank containing unlabeled cells (*p = 0*.*001, N = 4*) (**Figures 3A, B**). Consistent with this enhancement in R_1_, a significant increase in T_1_ weighted signal intensity could also be observed in the flank containing phage-labeled cells compared to the contralateral flank (49 ± 18 %, *p = 0*.*02, N = 4*) (**Figure 3C)**. This increase was also significant relative to the T_1_ weighted intensity measured in surrounding tissue background (43 ± 7 %, *p = 0*.*001, N = 4*) (**Figure S6**).

**Figure 3.**
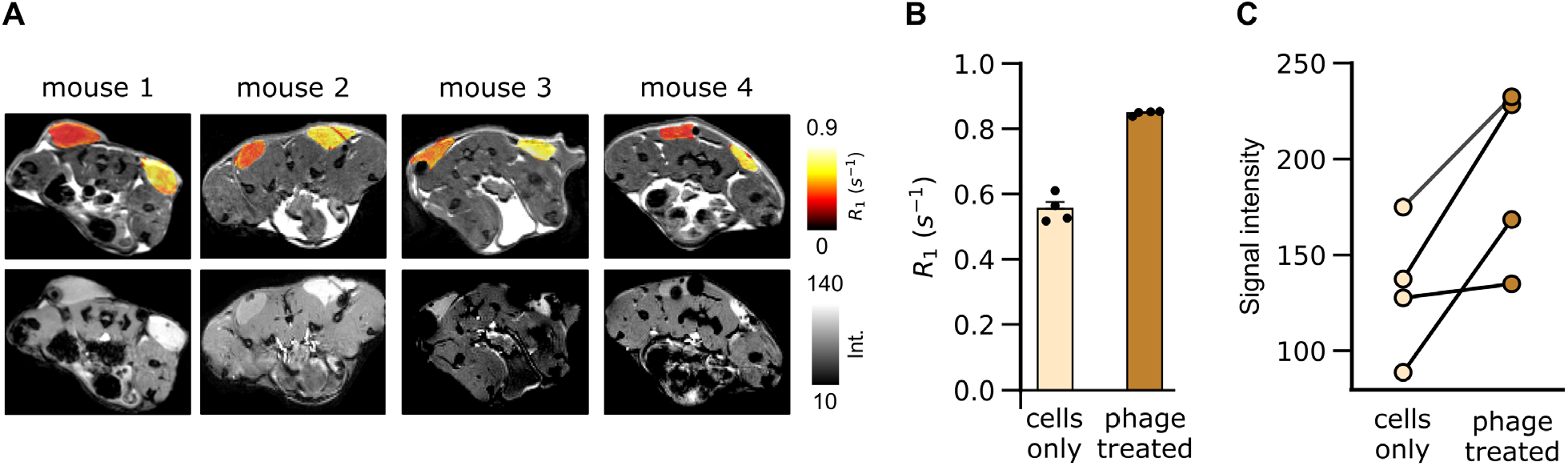
MR imaging of phage-labeled *V. cholerae* cells *in vivo*. **(A)** R_1_ maps (upper panel) and T_1_ weighted images (lower panel) of axial sections through the hind flanks of *N = 4* mice. The right flank of each animal was injected with phage-labeled *V. cholerae* cells, while the contralateral flank harbors phage-free *V. cholerae*. R_1_ maps are displayed as linear 8-bit heatmaps and overlaid on a T_2_ weighted anatomical image, thresholded for visual clarity. **(B)** R_1_ values and **(C)** T_1_ weighted signal intensities measured in each hind flank of the mice shown in (A). Imaging was performed on mice placed in a supine position in a vertical-bore 7 Tesla MRI scanner. Lines are drawn to connect paired values measured for bilateral flanks in the same subject. Error bars represent standard error of mean from *N = 4*

## CONCLUSIONS

In this work, we developed and applied a genetically engineered phage nanomaterial to image bacteria in a targeted fashion with MRI. While the current study demonstrated MR imaging of *E. coli* (F^+^) and *V. cholerae* cells, DP1-phage may in principle be engineered to noninvasively monitor different bacterial targets by tuning one of the four phage coat proteins (keeping pVIII for DP1 fusion) to incorporate receptor-binding proteins from native phages with tropism towards the desired bacterial strain^30^. The present work also established preliminary proof-of-principle for *in vivo* imaging of bacteria pre-labeled with DP1- phage, which may be useful for cell tracking applications involving the use of bacteria as live-cell therapeutics, especially if genetic modification of the bacterial strain to express reporter genes for noninvasive imaging^31–33^ is not desirable or possible. *In situ* detection of pathogenic or commensal microorganisms in their animal host would however require DP1-phage to be delivered *in vivo*. To this end, careful optimization of injected dose, injection route, frequency, and immunogenicity will be required for each bacterial type and anatomical site targeted with the phage. These efforts will be facilitated by a large body of literature addressing the delivery, biodistribution, and safety of filamentous phages in animal models ranging from mice to non-human primates^34–42^. In this regard, the ability to engineer phage variants capable of crossing vascular barriers (including the blood brain barrier^43–46^) could prove especially useful for imaging bacterial targets in hard-to-access locations such as the central nervous system.

Under our experimental conditions, we were able to detect 10^6^-10^7^ cells per voxel, which is comparable to net bacterial loads detected with previous phage-based sensors, albeit without the unique advantages of MRI; as well as using antibiotic-conjugated probes that are compatible with MRI but lack the cell-type specificity and tunability of the phage scaffold (**Table S3**). While contrast agents based on iron oxide nanoparticles have been reported to detect even fewer numbers of cells *in vivo* (on the order of 10^3^ per voxel), key drawbacks include the generation of negative contrast and limited engineering latitude compared to the highly tunable phage architecture. In the future, we anticipate that the sensitivity of DP1- phage could be further improved by engineering DP1 (for example, using phage display) to bind manganese with sub-micromolar affinity or fusing tandem repeats of DP1 to pVIII to increase manganese occupancy. In addition, theoretical considerations indicate that phage relaxivity should increase at lower fields (due to slowing down of molecular tumbling), which could also contribute to enhancing contrast at clinically-relevant field strengths (*e*.*g*., 1.5 Tesla), albeit with a tradeoff in overall signal intensity. In summary, as a phage-based molecular probe for visualizing targeted bacterial strains with MRI, we envision a synergy in the development of DP1-phage technology with rapidly evolving advances in emerging biotherapeutic platforms involving the use of live microorganisms as well as bacteriophage formulations for noninvasive diagnostics and treatment of diseases like cancer^46–52^, inflammatory conditions^53,54^, metabolic disorders^55,56^, and antibiotic-resistant infections^57–61^.

## METHODS

### Materials and reagents

Zinpyr-1, a fluorescent dye for assaying free manganese was purchased from Adipogen Corporation (San Diego, CA). *Pseudomonas aeruginosa* (Schroeter) Migula cells were purchased from ATCC (Manassas, VA, USA). *Vibrio cholerae* 0395 bacterial cells was a kind gift from Prof. Michael J. Mahan (UC, Santa Barbara). *E. coli* Nissle 1917 cells were harvested from a commercial probiotic formulation known as Mutaflor®. *E. coli* MG1655 cells were a kind gift from Prof. Charles M. Schroeder (University of Illinois at Urbana-Champaign). Reagents for Gibson assembly and mutagenesis as well as *E. coli* BL21 and *E. coli* ER2738 cells were purchased from New England Biolabs (Ipswich, MA). Oligonucleotide primers and gBlocks™ were obtained from Integrated DNA Technologies (Coralville, IA). Peptides for zinpyr-1 competition binding assay were synthesized by GenScript (Piscataway, NJ). Reagents for making competent *E*.*coli* cells were obtained from Zymo Research (Irvine, CA). C57BL/6J mice were purchased from The Jackson Laboratory (Bar Harbor, ME). Sterile hypodermic syringes (16G) were purchased from Air-Tite (North Adams, MA). Depilatory cream was obtained from Reckitt (Parsippany, NJ). Isoflurane anesthetic and activated charcoal (to scavenge residual isoflurane) were respectively purchased from Akorn Pharmaceuticals (Lake Forest, IL) and Henry Schein (Melville, NY). All reagents for protein electrophoresis and Western blotting were purchased from Biorad (Hercules, CA). Primary antibody (anti-M13 pVIII, #ab9225) was purchased from Abcam (Waltham, MA) and secondary antibody (HRP-conjugated rabbit anti-Mouse IgG, #61-6520) was purchased from Thermo Fisher. All other chemicals and reagents were purchased from MilliporeSigma (St. Louis, MO) or Thermo Fisher Scientific (Waltham, MA) and were of molecular biology grade.

### Fluorescence assay for manganese binding

Stock solutions of MnCl_2_ (1 mM) were prepared by dissolving the hydrated salt in deionized water. Stock solution of zinpyr-1 (1 mM) was prepared by dissolving the dye in dimethyl sulfoxide (DMSO) and stored in light-tight containers. Stock solutions of full-length (DEHGTAVM) and truncated DP1 peptides (DEHGTA) were prepared in deionized water. Manganese binding assays were performed in 300 μL reaction volumes comprising 4 μM zinpyr-1, 25 μM MnCl_2_, and peptide concentrations in the range of 0 - 200 μM. The mixture was incubated briefly at room temperature before measuring fluorescence with a microplate reader (Tecan Spark M200) by exciting zinpyr-1 with blue light (450 nm) and recording the emission spectrum from 485 to 630 nm. The monochromator slit width was set at 20 nm.

### Molecular biology

Phages were produced by sequentially transforming *E. coli* with 2 plasmids – a helper plasmid that supplies all 9 phage packaging proteins (pI- pIX) and a phagemid vector containing DNA sequences that can be packaged inside the phage capsid. The helper plasmid (plasmid #120346, HP17_KO7) was purchased from Addgene (Watertown, MA, USA) and modified by cloning the DP1 sequence at the N terminus of the pVIII gene downstream of the native signal peptide. The phagemid vector was constructed by amplifying an M13 origin of replication from pScaf (Addgene plasmid #111401) and sub-cloning the amplicon by Gibson assembly between the λ T0 and rrnB1 terminator sites in a GFP-expressing pQE80L expression vector constructed in-house. To design DP1-CTX phage, a gBlock™ encoding the receptor binding protein from *V. cholerae* CTXΦ phage was amplified by PCR and sub-cloned (by Gibson assembly) into the helper plasmid, swapping out the pilus-binding region (amino acids 18-236) of native pIII, but retaining the signaling sequence (amino acids 1-18). All genetic modifications were verified by Sanger sequencing (Genewiz, South Plainfield, NJ) and individual plasmids were propagated by transformation in competent *E. coli* NEB10 cells. See Table S1 for sequences.

### Phage purification

Four distinct types of phages were engineered in this work – wild type M13, DP1-M13, and DP1-CTX phage. To form phages, *E. coli* NEB10 cells were sequentially transformed first with the phagemid vector and then the helper packaging plasmid. Doubly transformed colonies were selected on LB-agar plates supplemented with ampicillin (100 μg/mL) and kanamycin (50 μg/mL). A single colony was used to generate a liquid culture by inoculation in 5 mL 2xYT medium (supplemented with kanamycin and ampicillin as before) in a 37 °C orbital shaker (set to 200 r.p.m). The overnight inoculum was sub-cultured at 1:100 dilution (v/v) into 20 mL fresh 2xYT medium (with antibiotics added as above) and grown for another 3-4 hours at 37 °C. This step yielded exponentially growing cells, which were then inoculated in 1 L fresh LB media in a 4 L flask and cultured as before. After overnight growth, the cells were harvested by centrifugation at 7,000 x g for 15 minutes. The supernatant (∼ 0.8 L) was removed and incubated with 0.2 L 5x precipitation buffer (2.5 M NaCl, 20 % PEG-8000 w/v). Following overnight incubation, phages were precipitated by centrifuging the PEG-treated supernatant at 12,000 x g for 15 minutes. The supernatant was decanted, and the phage pellet was resuspended in 40 mL HEPES buffer (20 mM, pH 7.0). Cell debris was removed by centrifuging the phage resuspension at 15,000 x g for 15 minutes. The supernatant was mixed one more time with 10 mL precipitation buffer and chilled on ice for 15-60 minutes before precipitating phage particles by centrifugation at 12,000 x g for 15 minutes. The final pellet was resuspended in 10 mL HEPES buffer (20 mM, pH 7.0). All precipitation steps were carried out at 4 °C. Phages were verified by Western blotting and phage titer was quantified using absorbance spectroscopy as follows: 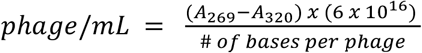 Here, *AA*_269_ and *AA*_320_ are the absorbance values of the phage suspension at 269 and 320 nm respectively, and the number of bases is 5153 corresponding to the size of the phagemid vector. Phages prepared in this way were routinely stored at 4 °C without any noticeable loss of target cell binding (as assessed by MRI).

### Western blotting

Phages were diluted in HEPES buffer (20 mM, pH 7.0) to ∼ 24 μM (based on pVIII concentration), mixed 1:1 (v/v) with 2x Laemmli Buffer supplemented with 5% 2-mercaptoethanol, and denatured by heating at 95 °C for 10 minutes. Approximately 20 μL of the denatured phage was loaded in a 4-20 % gradient gel (Mini-PROTEAN® TGX Stain-Free gel, Biorad) and separated by electrophoresis at 120 V. Proteins were transferred to a 0.45 μM PVDF membrane using a Trans-Blot Turbo Transfer System (Biorad). In preparation for Western blotting, blocking buffer was made by dissolving 5% non-fat dry milk in tris-buffer saline supplemented with 0.05 % Tween-20 (TBS-T buffer). Antibody dilutions were prepared in blocking buffer at 1:1000 (v/v) for the primary antibody (anti-M13 pVIII) and 1:2000 (v/v) for the secondary antibody (HRP-conjugated rabbit anti-Mouse IgG). After transfer was complete, the PVDF membrane was washed several times with deionized water to remove excess transfer buffer, and subsequently incubated in blocking buffer at room temperature with orbital shaking (200 r.p.m.). After 30 minutes, the membrane was transferred to the primary antibody solution and incubated at 4 °C with gentle shaking (200 r.p.m.). Following overnight incubation, primary antibody was removed by washing three times with TBS-T and the membrane was then incubated with secondary antibody at room temperature for 1 hour (200 r.p.m.). Finally, the membrane was washed three times in TBS-T, treated with a chemiluminescent substrate (Clarity Western ECL Substrate, Bio-Rad), and imaged in a gel imager.

### Analytical size exclusion chromatography

PEG-precipitated phage particles (0.5 mL) were injected in a Superdex 200 size exclusion column and washed with at least two column volumes of HEPES buffer at a flowrate of 0.5 mL/min. Elution volume of phage was monitored by inline recording of absorbance at 280 nm.

### Metal loading of purified phage particles

To load phages with Mn^2+^, purified phage particles were incubated with manganese chloride solution at 5:1 (Mn^2+^ to DP1) molar stoichiometry, mixed briefly by vortexing (∼ 5 seconds), and then placed in a rotator (∼ 15 r.p.m.) for approximately 15 minutes. Molarity calculations assumed 2172 pVIII molecules were present in an individual phage particle. After mixing, the phage-manganese mixture was treated with phage precipitation buffer and chilled on ice for 30 minutes. Phages were subsequently purified by 3 rounds of PEG-precipitation and centrifugation (17,000 x g, 10 minutes at 4 °C), resuspended in 200-300 μL HEPES buffer (20 mM, pH 7.0), and quantified by measuring absorbance as described above. Phages were always freshly loaded with manganese before each experiment. To test binding specificity, the Mn^2+^-treated phage was resuspended in HEPES buffer supplemented with divalent metals at physiologically relevant (extracellular) concentrations: 1.8 mM Ca^2+^, 1 mM Mg^2+^, or 10 μM Zn^2+^. Phages were subsequently purified by 2 rounds of PEG-precipitation, resuspended in 200-300 μL HEPES buffer (without added metals), quantified, and characterized by relaxometric titration to measure T_1_ relaxation rates (R_1_). In some instances, purified phage was resuspended in mouse serum (Thermo Fisher) or Dulbecco’s Modified Eagle Medium (DMEM) instead of HEPES and characterized as above to measure R_1_.

### Inductively coupled plasma-mass spectrometry (ICP-MS)

ICP-MS experiments were performed using plasticware soaked in 10% nitric acid overnight and rinsed three times with deionized water. The use of glassware was avoided to prevent contamination due to leaching of impurities from glass. Mn^2+^ free and bound phages were prepared as described above and dissolved in 5% nitric acid. Diluted phage samples were then analyzed for manganese content using an Agilent 7900 ICP-MS (Agilent Technologies, Santa Clara, CA) at the UC Santa Barbara Bren School of Environmental Science and Management.

### Preparation of cells for MRI

Starter cultures of bacterial cells (*E. coli* ER2738, *E. coli* Nissle, *E. coli* BL21, *V. cholerae*, and *P. aeruginosa*) were grown in 5 mL lysogeny broth overnight at 37 °C. For *E. coli* ER2738 cells, the growth medium was supplemented with tetracycline (10 µg/mL) to maintain episomal expression of the F-pilus. The pre-inoculum was added to fresh lysogeny broth at 1:100 dilution (v/v) and grown for 12-16 h at 37 °C. We quantified cell density by measuring optical density at 600 nm. 10 mL of the overnight culture was incubated with manganese-loaded phage particles at a phage-to-cell stoichiometry of 50:1, mixed by inverting for 5 minutes, and centrifuged at 3,000 x g for 10 min. The supernatant was decanted, and the cells were resuspended in 1 mL HEPES buffer. The cells were washed one more time and finally resuspended in 150 μL HEPES in MRI-compatible plastic tubes. The resuspension was centrifuged at 3,000 x g for 10 minutes to form pellets for imaging with MRI. Mammalian cells (Jurkat and RAW 264.7) were cultured in RPMI medium supplemented with 10 % fetal bovine serum, 100 U/mL penicillin, and 100 μg/mL streptomycin, at 37 ºC, 5 % CO_2_ in a humidified chamber. Jurkat cells were grown as suspension cultures and 10 mL of the culture was incubated with phage as described before. Adherent RAW264.7 cells were grown till 70-90 % confluency, detached by treating with trypsin-EDTA, and incubated with phage as described above. Mammalian cells were pelleted by centrifuging at 350 x g for 10 minutes at 4 °C, the lower speeds (compared to bacteria) necessary to prevent cell lysis. The supernatant was decanted; and the cell pellet was resuspended in 1 mL HEPES buffer. The wash step was repeated one more time before resuspending the cells in a 150 μL HEPES. The cells were finally centrifuged at 500 x g for 10 minutes to form pellets for analysis by MRI.

### *In vitro* MRI

In preparation for MRI, plastic tubes containing cells or phage suspensions were placed in water-filled agarose (1% w/v) moulds in a 3D printed MRI phantom designed to minimize magnetic susceptibility differences between the samples and the surrounding environment. All *in vitro* MRI experiments were performed in a Bruker 7 Tesla vertical bore MRI scanner using a 66 mm diameter transceiver coil. To measure T_1_, multiple images were acquired using a rapid acquisition with relaxation enhancement (RARE) pulse sequence with the following parameters: echo time, T_E_: 12.6 ms, RARE factor: 4, field of view: 4.7 cm x 4.7 cm, matrix size: 128 × 128, slice thickness: 1.5 – 2 mm, and variable repetition times, T_R_: 173.5, 305.9, 458.6, 638.8, 858.7, 1141.1, 1536.1, 2198.4, and 5000 ms.

### Subcutaneous cell implantation and *in vivo* MRI

*V. cholerae* cells were labeled with CTX-DP1 dual functionalized phage as described above, resuspended in 165 μL Matrigel® at 4 °C, and centrifuged at 3,000 x g for 10 minutes. After decanting the supernatant, the cells were stirred into a slurry and carefully aspirated into a pre-chilled 16G syringe with the headspace pre-filled with sterile saline solution. In preparation for *in vivo* injection, the syringe was warmed at room temperature for ∼ 10 minutes. Female C57BL/6J mice between 5- 7 weeks of age were anesthetized with 2.5 % isoflurane and secured in an animal cradle with anesthetic gas (1 – 2 % isoflurane in medical oxygen) continuously delivered through a nosecone. Approximately 100 μL cell slurry was injected subcutaneously in one flank, while identically prepared phage-free cells were implanted in the contralateral flank of the same animal to serve as control. Imaging was performed in a Bruker 7 Tesla vertical bore MRI scanner using a 40 mm diameter transceiver coil. Throughout the imaging session, body temperature was maintained at ∼ 37 °C by circulating warm air with feedback-driven temperature control. Respiration rate (beats per minute) and body temperature were continuously monitored with a pneumatic transducer (Biopac Systems, Goleta, CA) and fiber optic rectal probe (OpSens, Quebec, Canada) respectively. Locations of the implanted cells in each flank were determined from a multi-slice anatomical (T_2_ weighted) scan acquired with the following parameters: T_R_: 2.5 s, T_E_: 40 ms, RARE factor: 4, number of averages: 5, matrix size: 256 × 256, field of view: 3.13 cm x 3.13 cm, slice thickness: 1 mm, in-plane resolution: 122 × 122 μm^2^. T_1_ weighted images were subsequently acquired using a FLASH sequence with the following parameters: T_R_: 83.7 ms, T_E_: 3 ms, flip angle: 30 °C, averages: 10, matrix size: 128 × 128, field of view: 3.13 cm x 3.13 cm, slice thickness: 1 mm, in-plane resolution: 244 × 244 μm^2^. Finally, to measure T_1_, images were acquired at multiple repetition times over the same field of view as before using a RARE spin echo sequence with identical parameters as described above for *in vitro* MRI. All animal procedures were approved by the Institutional Animal Care and Use Committee at UC Santa Barbara (Protocol #946).

### Image processing and analysis

ImageJ (NIH) was used to measure average signal intensities in manually drawn regions of interest encompassing cell samples. To estimate T_1_ relaxation rates (R_1_), the change in signal intensity as a function of repetition time (T_R_) was fitted to the equation 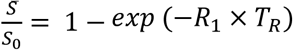, which describes the growth of magnetization by longitudinal relaxation following application of a radiofrequency pulse. Molar relaxivity (r_1_) was derived by linear regression of R_1_ *vs*. the concentration of pVIII protein (estimated from the phage titer). To generate voxel-wise R_1_ maps, the signal intensity in each voxel (within a region of interest) was fitted to the exponential growth equation described above. The resulting images were smoothed with a 2 pixel wide median filter and displayed on a linear 8-bit color scale. For *in vivo* images, voxel-wise R_1_ maps were overlaid on a T_2_ weighted anatomical image in the same plane. Least squares regression and voxel-wise mapping were implemented in Matlab (v. 2018a).

## Data analysis

All measurements are reported as mean ± standard error based on experiments conducted using *N ≥ 3* independent replicates. For least squares regression, quality of model fits was determined from the regression coefficient. Pairwise comparisons were performed using paired or unpaired Student’s t-test with significance level set at 0.05.

## Author Contributions

AM and IAC conceived the study. HFO, REB, BX, HP, IAC, and AM designed experiments. REB, HFO, and BX performed all experiments with help from XF and EL who assisted with bulk phage purification, gene sequencing, phage quantification, and relaxometric assays. REB, HFO, BX, and AM performed all data analysis. AM wrote the manuscript with inputs from all authors. AM and IAC supervised the research.

## Acknowledgements

We thank members of the Mukherjee and Chen labs for helpful discussions. We thank Prof. Mikhail Shapiro (Caltech) for several insightful discussions on MRI-based bacterial imaging MRI. Dr. Jerry Hu (UC, Santa Barbara) is gratefully acknowledged for assistance with setting up the MRI workflow. This work was supported by the California NanoSystems Institute (University of California, Santa Barbara) and a National Institutes of Health R35 Maximizing Investigators’ Research Award (5R35GM133530; AM), NIGMS (DP2 GM123457; IAC), and the Camille Dreyfus Teacher-Scholar Awards Program (IAC). HFO gratefully acknowledges support from the Errett Fisher Foundation. All MRI experiments were performed at the Materials Research Laboratory (MRL) at UC, Santa Barbara. The MRL Shared Experimental Facilities are supported by the MRSEC Program of the NSF under Award No. DMR 1720256; a member of the NSF-funded Materials Research Facilities Network.

## SUPPORTING INFORMATION

**Figure S1.**
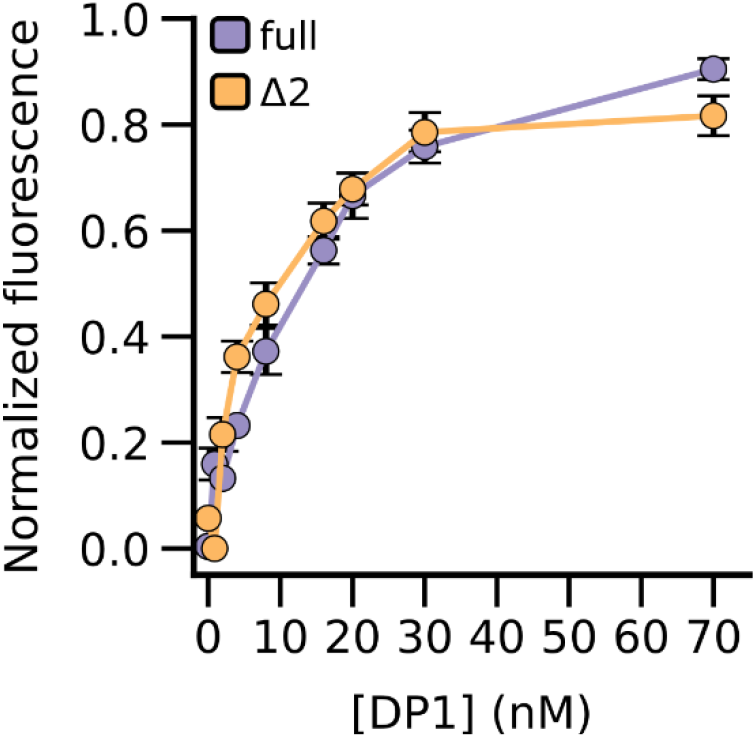
Fluorescence assay for manganese binding to DP1. Binding was assessed by measuring fluorescence recovery of quenched zinpyr-1 dye (4 μM) while the full-length (purple) or truncated (orange) form of DP1 was titrated from 0-70 μM. Initial MnCl_2_ concentration in the reaction volume was 25 μM. Fluorescence was measured by exciting the reaction volume at 450 nm and integrating the emission spectrum from 485 to 630 nm. Values are normalized to peak fluorescence measured at saturating DP1 (200 μM). Error bars represent standard error of mean of *N = 3* replicates.

**Figure S2.**
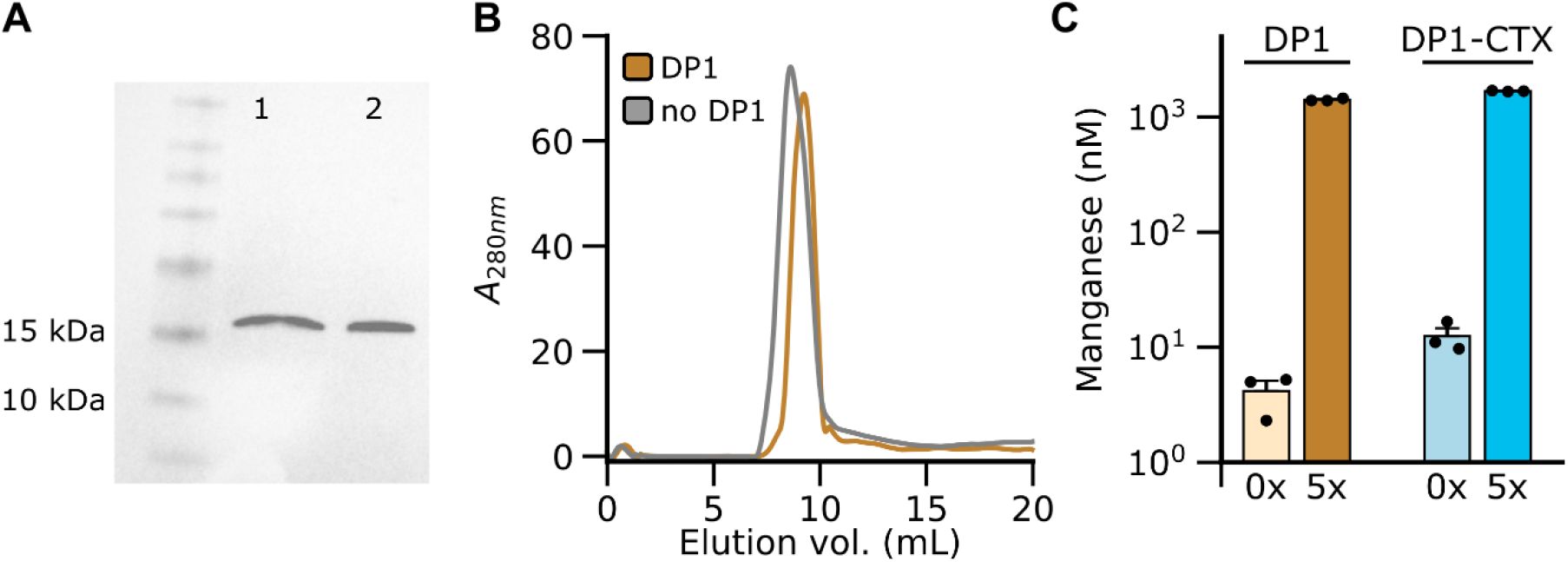
Characterization of DP1-phage. **(A)** Formation of DP1-phage (lane 2) was verified with Western blotting using an antibody targeted against pVIII. Lane 1 contains identically purified phage lacking DP1. **(B)** Phage formation was also assayed using analytical size exclusion chromatography. DP1-phage was found to elute at nearly the same volume as phage lacking DP1. **(C)** Manganese loading in DP1- and DP1-CTX phages was confirmed with inductively coupled plasma mass spectrometry (ICP-MS). In these experiments, the phage titer (assayed with absorbance spectroscopy) corresponds to ∼ 3 μM pVIII. Bar labels (0x and 5x) indicate the relative amount of manganese that was initially incubated with the phage. Accordingly, 47.3 ± 0.9 % and 54.3 ± 4.7 % of the phage capsid was loaded with manganese respectively for DP1- and DP1-CTX phages, respectively. Error bars represent standard error of mean from N = 3 replicates.

**Figure S3.**
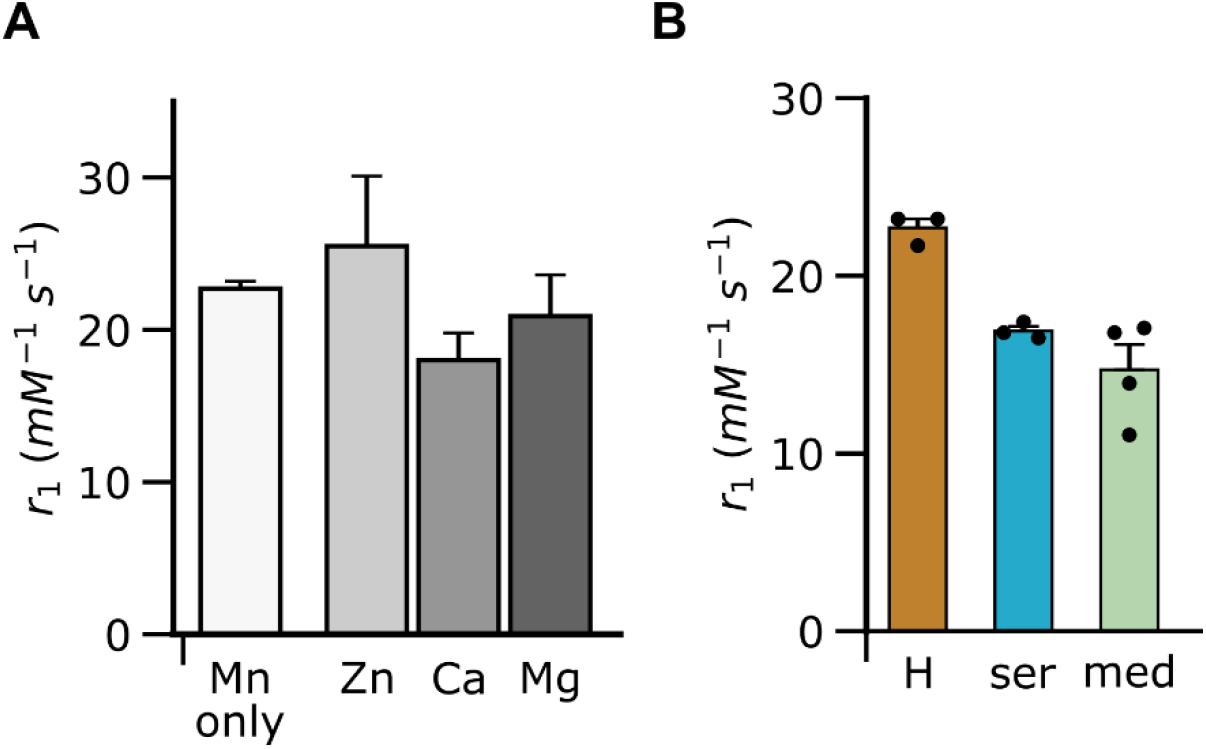
Relaxometric characterization of DP1-phage in physiological conditions. **(A)** Specificity of DP1-phage towards manganese was assessed by measuring relaxivity (r_1_) following incubation of phage (pre-loaded with Mn^2+^) in HEPES buffer containing other (non-paramagnetic) divalent metal ions supplemented at biologically relevant extracellular concentrations: 10 uM (Zn^2+^), 1.8 mM (Ca^2+^), and 1 mM (Mg^2+^). No substantial change in r_1_ was detected under these conditions (*p > 0*.*6, N = 4*). **(B)** Relaxivity of DP1-phage in mouse serum and cell culture medium showed a modest decrease compared to aqueous buffer. H: HEPES buffer, ser: mouse serum, med: Dulbecco’s Modified Eagle Medium (DMEM). Error bars represent standard error of mean from N ≥ 3 replicates.

**Figure S4.**
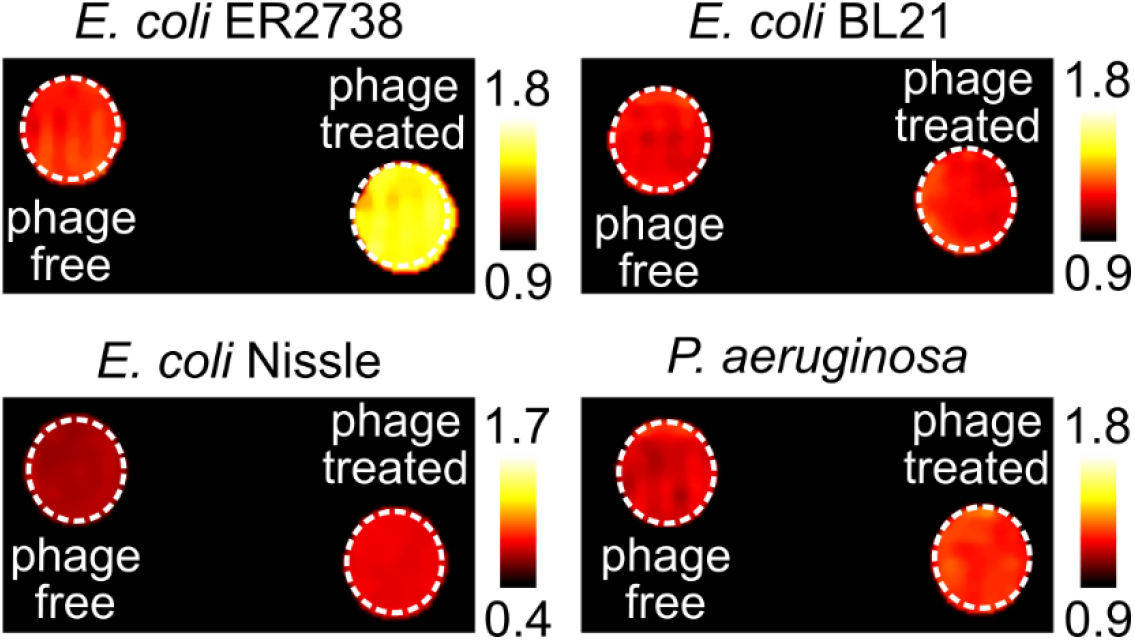
MRI-based detection of *E. coli* (F^+^) using DP1-phage. R_1_ maps of axial sections through bacterial cell pellets contained in plastic tubes placed inside a 1% (w/v) agarose phantom to minimize susceptibility differences. Unlabeled (*i*.*e*., phage-free) cells were imaged side by side with cells treated with Mn^2+^-loaded DP1-phage. A detectable signal change was observed only in case of the target *E. coli* F(^+^) strain, ER2738. R_1_ maps were generated by manually masking the region of interest (ROI) and fitting the signal intensity in each voxel within this ROI to an exponential equation describing T_1_ relaxation. The images were smoothed with a 2-pixel median filter and displayed on a linear 8-bit heatmap by thresholding at minimum and maximum values of R_1_ as depicted in the adjoining color bars (units are s^-1^).

**Figure S5.**
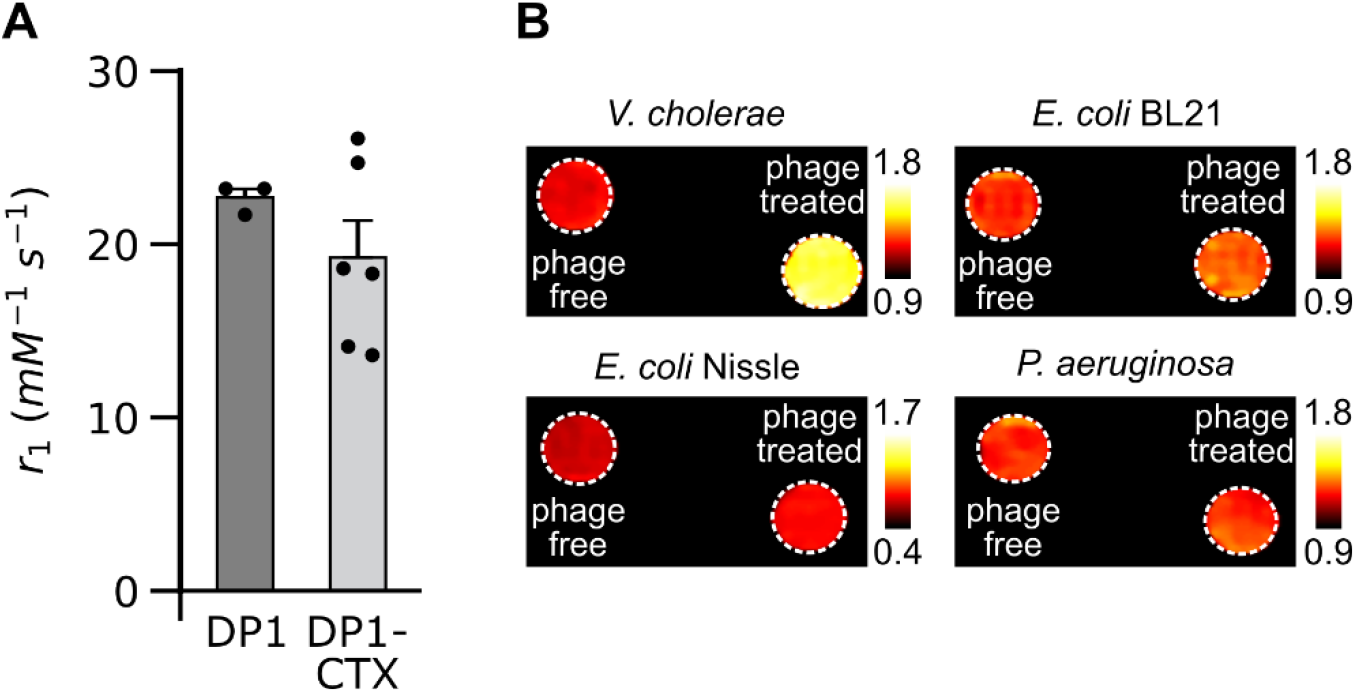
MRI-based detection of *V. cholerae* using DP1-CTX phage. **(A)** DP1-CTX phage shows comparable relaxivity as DP1-phage. Error bars represent standard error of mean from N ≥ 3 replicates. **(B)** R_1_ maps of axial sections through bacterial cell pellets contained in plastic tubes placed inside a 1% (w/v) agarose phantom to minimize susceptibility differences. Unlabeled (*i*.*e*., phage-free) cells were imaged side by side with cells treated with Mn^2+^-loaded DP1-phage. A detectable signal change was observed only in case of the target cells, *i*.*e*., *V. cholerae*. R_1_ maps were generated by masking the ROI and fitting the signal intensity in each voxel within this ROI to an exponential equation describing T_1_ relaxation. The images were smoothed with a 2-pixel median filter and displayed on a linear 8-bit heatmap by thresholding at minimum and maximum values of R_1_ as depicted in the adjoining color bars (units are s^-1^).

**Figure S6.**
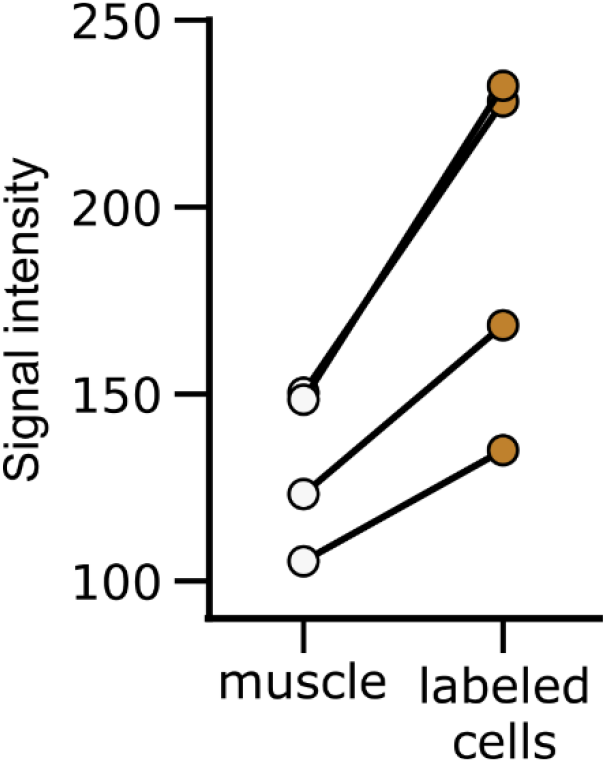
*In vivo* imaging of phage-labeled *V. cholerae*. T_1_ weighted signal intensity measured in muscle of the hind flanks harboring subcutaneously implanted CTX-phage-labeled *V. cholerae*. The flank containing phage-labeled cells showed a significant increase in signal intensity compared to background tissue (*p = 0*.*02, N = 4, paired t-test*).

**Table S1.**
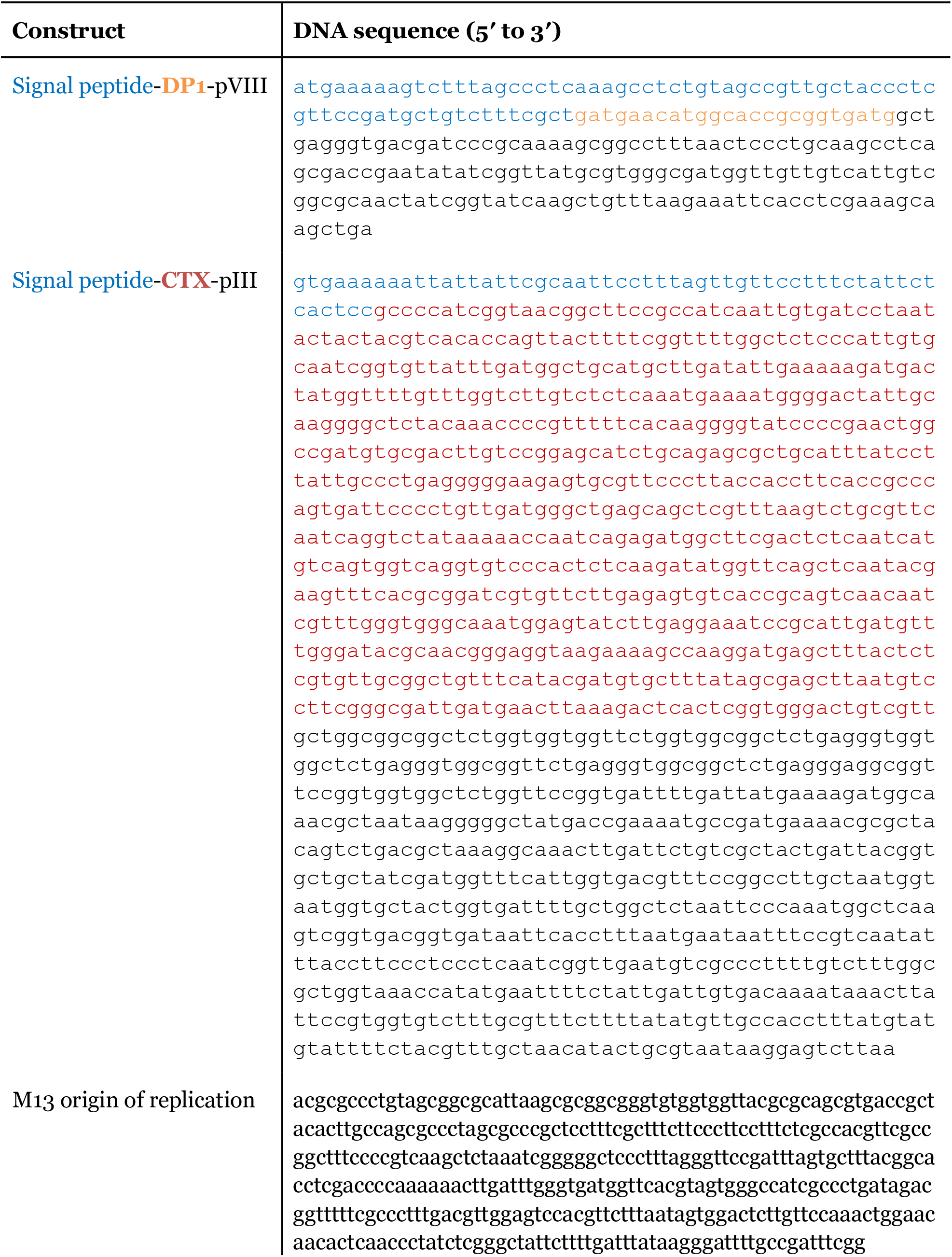
Genetic constructs engineered in this work.

**Table S2.** Phage-based probes for bacterial imaging.

| Probe | Modality | Phage scaffold | Detection (cells/mL) | Bacterial targets | Potential limitations |
| --- | --- | --- | --- | --- | --- |
| Single-walled nanotubes<sup>1</sup> | near infrared | M13 | 10<sup>8</sup>-10<sup>9</sup> | *E. coli* (F<sup>+</sup>), *S. aureus* | limited depth penetration & spatial resolution |
| Alexa750<sup>2</sup> | near infrared | M13 | 10<sup>9</sup> | *E. coli* (F<sup>+</sup>), *S. aureus* | limited depth penetration & spatial resolution |
| <sup>99m</sup>Tc<sup>3</sup> | positron emission | M13 | 5 x 10<sup>8</sup> | *E. coli* 25922, *E. coli* 2537, *S. aureus* | needs ionizing radiation, limited spatial resolution |

**Table S3.**
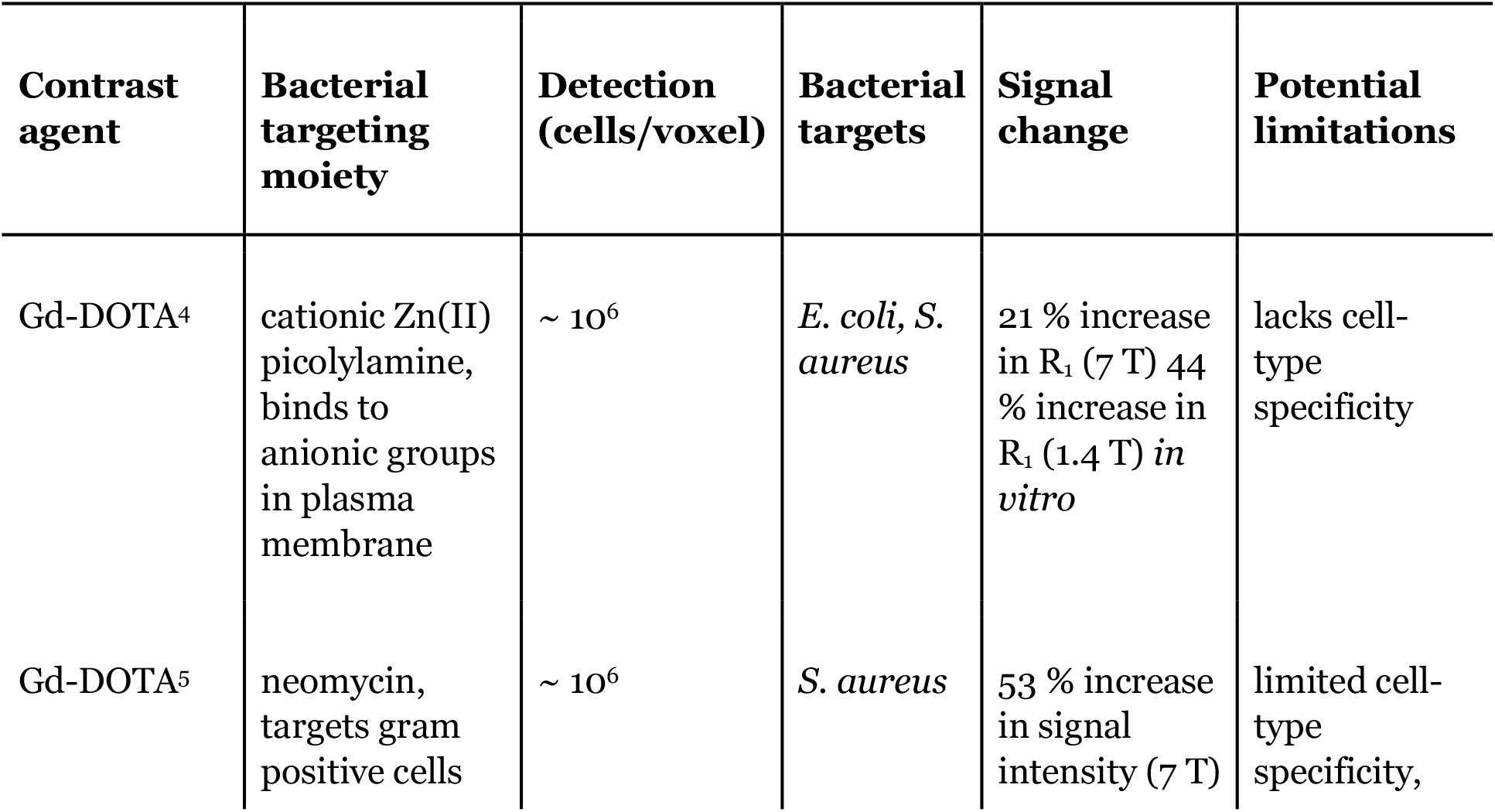
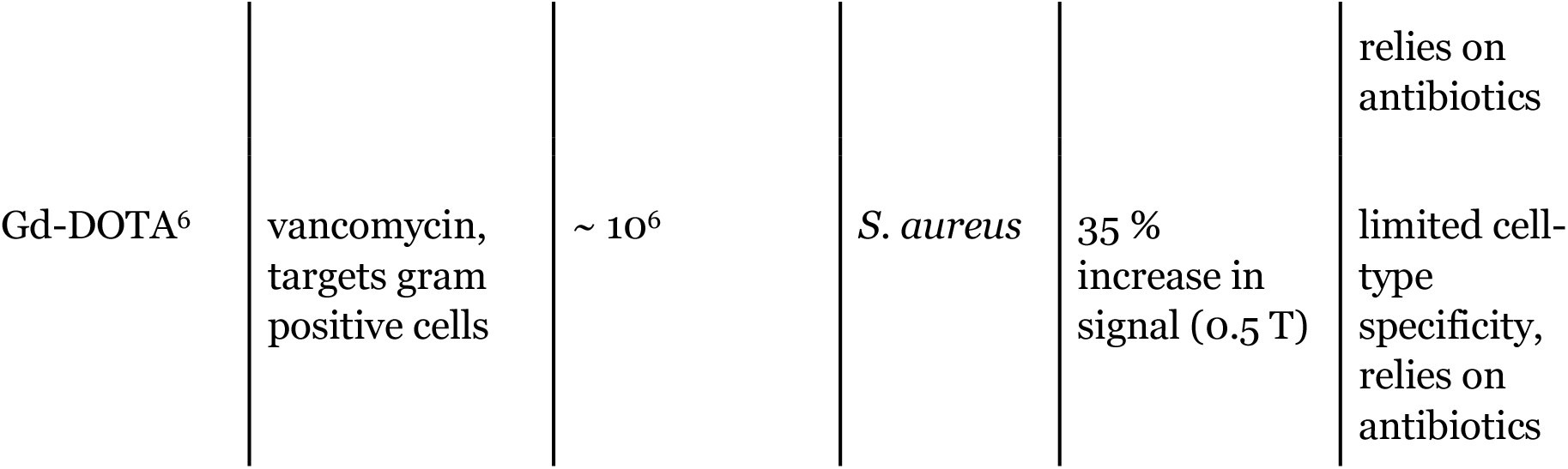
T_1_ weighted MRI probes for bacterial imaging.

